# PPIFold: a tool for analysis of Protein-Protein Interaction from AlphaPullDown

**DOI:** 10.1101/2025.01.08.631489

**Authors:** Quentin Rouger, Emmanuel Giudice, Damien F. Meyer, Kévin Macé

**Affiliations:** Univ. Rennes, CNRS, Institut de Génétique et Développement de Rennes (IGDR) - UMR6290, 35000 Rennes, France; CIRAD, UMR ASTRE, 97170 Petit-Bourg, Guadeloupe, France; ASTRE, University Montpellier, CIRAD, INRAE, Montpellier, France

## Abstract

Protein structure and protein-protein interaction (PPI) predictions based on coevolution have transformed structural biology, but managing pre-processing and post-processing can be complex and time-consuming, making these tools less accessible. Here, we introduce PPIFold, a pipeline built on the AlphaPulldown Python package, designed to automate file handling and streamline the generation of outputs, facilitating the interpretation of PPI prediction results. The pipeline was validated on the bacterial Type 4 Secretion System nanomachine, demonstrating its effectiveness in simplifying PPI analysis and enhancing accessibility for researchers.

## 1. Introduction

Artificial intelligence has revolutionized the field of structural biology with software such as AlphaFold2 (*Jumper et al., Nature, 2021*) and RoseTTAFold (*Baker et al., Science, 2021*). In addition to predicting structure, these methods have proven very effective in predicting protein-protein interaction (PPI) (*Humphreys et al., Nature Microbiology, 2024*). In the context of high-throughput PPI prediction, the process often involves a series of redundant and automatable steps, along with essential verification stages to ensure interpretable and reliable results. The AlphaPulldown tool (*Yu et al., Bioinformatics, 2023*) exemplifies this approach but requires considerable manual intervention to achieve high-confidence predictions. To address these challenges, we present PPIFold, a Python-based tool designed to streamline PPI prediction by minimizing redundant steps and reducing the likelihood of inaccurate results. PPIFold is a more intuitive and specialized tool for protein-protein interaction (PPI) analysis, incorporating new interaction scoring metrics and automating the generation of detailed interaction evaluation reports. This enhanced functionality allows for a more comprehensive assessment of predicted PPIs, streamlining the interpretation process and providing deeper insights into interaction dynamics, making it a powerful tool for PPI-focused research.

PPIFold offers an accessible solution for researchers, particularly those without extensive bioinformatics expertise, enabling them to efficiently predict PPIs by eliminating unnecessary steps and rapidly generating the data and figures necessary for publication. The tool is built upon the AlphaPulldown package, which leverages AlphaFold Multimer (*Evans et al., Bioinformatics, 2022)* for extensive PPI screening across diverse protein combinations. Additionally, this pipeline allows for the separation of feature generation (handled by CPUs) from model generation (handled by GPUs), further optimizing the computational workflow.

## 2. Pipeline description

The PPIFold pipeline is designed for the identification or validation of protein-protein interactions (PPI), including homo-oligomers (*Fig. 1*). PPIFold is structured into three main modules. First, protein sequences undergo automatic cleaning and verification. Next, the pipeline predicts all potential PPIs using the AlphaPulldown tool. Finally, the predicted interactions are scored, and detailed figures are generated for the PPIs of interest, facilitating a comprehensive analysis and visualization of the results (*Supplementary Information*).

**Figure 1.**
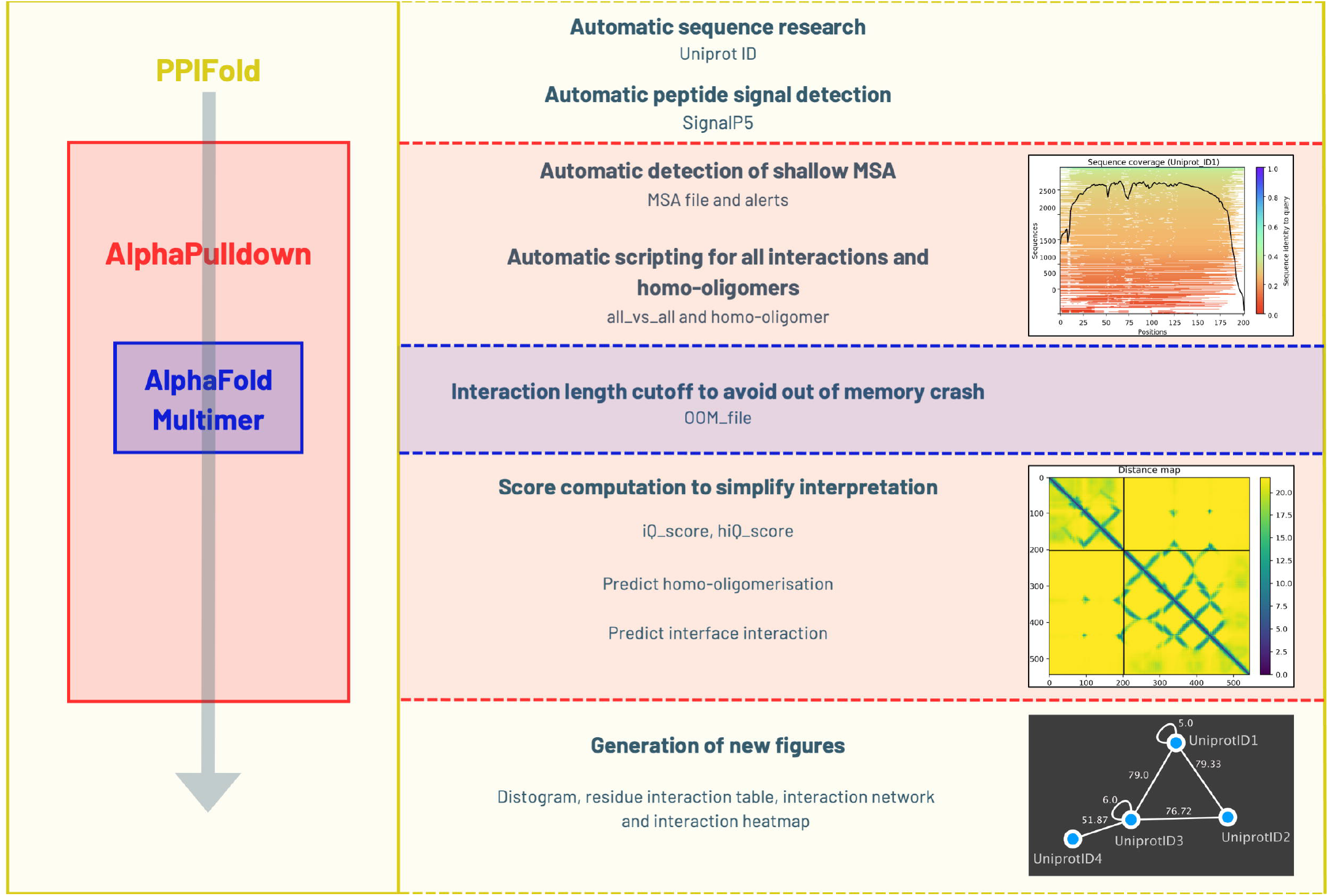
Depiction of the PPIFold pipeline. The PPIFold pipeline consists of three major modules. The first component involves sequence generation and verification, including the creation of scripts for AlphaPulldown. The second component focuses on model generation using AlphaPulldown, which provides initial scores for predicted protein-protein interactions. The final component involves calculating refined scores and generating various figures and tables for data visualization and interpretation.

### 2.A. Sequence and verification

Inputs sequences are analyzed using SignalP5 (*Almagro Armenteros et al., Nature Biotechnology, 2019*) to identify the presence of signal peptides characteristic of the organism, and if yes, the sequences are removed from the original protein sequences to obtain the mature protein structure post-translocation into the periplasm. Next, the feature generation and multiple sequence alignment (MSA) for all proteins are performed using mmseq2 or hmmer *via* AlphaPulldown package. Then PPIFold generates an MSA depth figure to assess the quality of the alignment, as a shallow MSA can negatively impact prediction accuracy, the alignment depth is adequate to capture co-evolutionary signals, which are crucial for predicting both intra- and inter-protein interactions. Proteins with poor MSA quality are recorded in a file named bad_MSA.txt, and predictions for these proteins are flagged as unreliable for validating or invalidating protein-protein interactions.

Additionally, PPIFold provides two .txt files of interaction prediction to do, all-against-all and homo-oligomers, tailored according to the available GPU memory (https://www.rbvi.ucsf.edu/chimerax/data/alphafold-jan2022/afspeed.html). Indeed, to prevent out-of-memory (OOM) errors, interactions too large to model are excluded, and these cases are listed in the OOM_int.txt file, allowing users to track interactions that cannot be processed due to memory constraints.

### 2.B. Structures prediction by AlphaPulldown

The custom mode and homo-oligomer functions from AlphaPulldown tool *via* AlphaFold Multimer are utilized to generate structural models and assign various scores to rank predicted protein-protein interactions (PPIs). For each interaction, five different models are generated and ranked based on their pLDDT scores (*Mariana et al., Bioinformatics, 2013*), with only the highest-ranked model undergoing further scoring. This step, which is the most time-intensive, is GPU-dependent.

### 2.C. Scores

Interaction within hetero-oligomeric models are evaluated using three distinct scores generated by AlphaPulldown tool: pi-score (*Malhotra et al., Nature communications, 2021*), iptm/ptm scores (*Gao et al., Nature communications, 2022*), and pDockQ (*Bryant et al., Nature communications, 2022*). On the other hand, interaction within homo-oligomeric models are assessed using two scores: pi-score and iptm/ptm. While each score provides valuable insights into a protein-protein interaction, relying on a single score is insufficient, moreover evaluating multiple scores can be difficult and time-consuming. To address this, we introduced the iQ-score, a weighted linear combination of these individual scores, with an emphasis on the pi-score. This iQ-score reduced false positives, especially since some proteins, such as small or membrane-associated proteins, may exhibit high scores across most tested interactions. The iQ-score incorporates cutoffs based on those of the individual scores from which it is derived, offering a more streamlined and reliable assessment of PPIs.

*iQ-score = ((pi-score+2,63)/5,26)*40+iptm_ptm*30+pDockQ*30*

*iQ-score >= 35,37*

*hiQ-score = (((*∑*pi-score)/n+2,63)/5,26)*60+iptm_ptm*40*

*hiQ-score >= 52,6*

These scores are calculated exclusively for the better model (higher pLDDT) and for interactions with a predicted alignment error (PAE) above a threshold of 10, which is directly influenced by the depth of the multiple sequence alignment (MSA). All results are sorted based on these cutoff scores, ensuring that only the most probable interactions are retained. Then, PPIFold does further analysis and figures generation, that includes a distogram figure, a residue interaction table, a heatmap and a protein interaction network figure.

### 2.D Figures generation

#### Distogram

The distogram figure provides a rapid and schematic visualization of the interface involved in the protein-protein interaction. It represents interactions into two proteins where the value on the left indicates the protein sequence length in amino acids. Points near the diagonal symmetry line, along with pixels in black squares, represent residues in contact within the same protein, while points outside this region denote residues in contact between different proteins. Colors correspond to the distance between residue pairs, measured in Ångströms, with dark blue points indicating shorter distances. This distogram is generated directly from the PDB file and is displayed only for protein-protein interactions that have passed the cutoff scores.

#### Residue interaction table

For each interaction, all residues in contact at the interface are listed in the table, providing a more detailed representation of the distogram. This additional information is particularly useful for designing mutations for wet lab validation. The table is generated exclusively for the top-ranked protein-protein interactions that meet the cutoff criteria.

#### Heatmap and interaction network

The iQ-score heatmap allows a comprehensive visualization of interaction scores for all proteins, highlighting those with either low or high average scores. The interaction network figure represents all proteins that have at least one interaction with other proteins within the system indicated by iQ-score. It also illustrates the homo-oligomerization of each protein indicated by hiQ-score. Thus, the interaction network provides a comprehensive overview of the protein-protein interactions, facilitating the generation of hypotheses regarding the system’s structure and function. By examining the network, researchers can explore functional relationships and quickly identify potential incompatibilities between interactions. These observations can be further supported by detailed analyses of interaction interfaces on the PDB models, such as when two proteins interact at the same interface area of a third partner. The scoring system serves as an indicator of interaction strength, offering a valuable tool for predicting the stability and likelihood of protein interactions within the system.

## 3. Conclusion

PPIFold builds on the strengths of AlphaPulldown and AlphaFold Multimer, automating and simplifying both pre- and post-processing steps for large-scale protein-protein interaction (PPI) predictions. By streamlining the workflow, PPIFold reduces manual intervention, making PPI analysis more efficient and accessible to a broader community of researchers. This pipeline accelerates the *in silico* exploration of PPIs, offering a fast and efficient tool for generating biological hypotheses, improving experimental design, and contributing to a deeper understanding of molecular interactions. Moreover, PPIFold stands out as an essential resource for researchers seeking to harness cutting-edge computational power, ultimately enabling groundbreaking discoveries in structural and systems biology. Looking ahead, PPIFold is designed with the flexibility to be adapted to AlphaFold3 or any future prediction software advanced.

## SUPPLEMENTARY INFORMATION & DATA

PPIFold tools were evaluated on the 11 proteins constituting the Type 4 Secretion System (T4SS) nanomachine to predict protein-protein interactions within this complex. T4SS R388 was chosen as a model system to benchmark PPIFold’s predictions against the experimentally resolved cryo-EM structure (*Macé et al., Nature, 2022*). Supplementary Figures 1–3 focus on the interactions between two specific proteins, O50331 (TrwJ) and O50333 (TrwI), while Supplementary Figures 4–5 present the entire interactome prediction as a heatmap and a network representation. Additionally, the complete T4SS R388 analysis results are provided in Supplementary Data 1 (all-vs-all interactions) and Supplementary Data 2 (homo- oligomerisation predictions).

## DATA AVAILABILITY

This pipeline with test T4SS dataset is available on Github (https://github.com/Qrouger/PPIFold)

## ACKNOWLEDGMENTS

The authors thank Alix Regnier for reviewing the code.

