## Supplementary Information for "PPIFold: a tool for analysis of Protein-Protein Interaction from AlphaPullDown"

**SUPPLEMENTARY FIGURE 1**

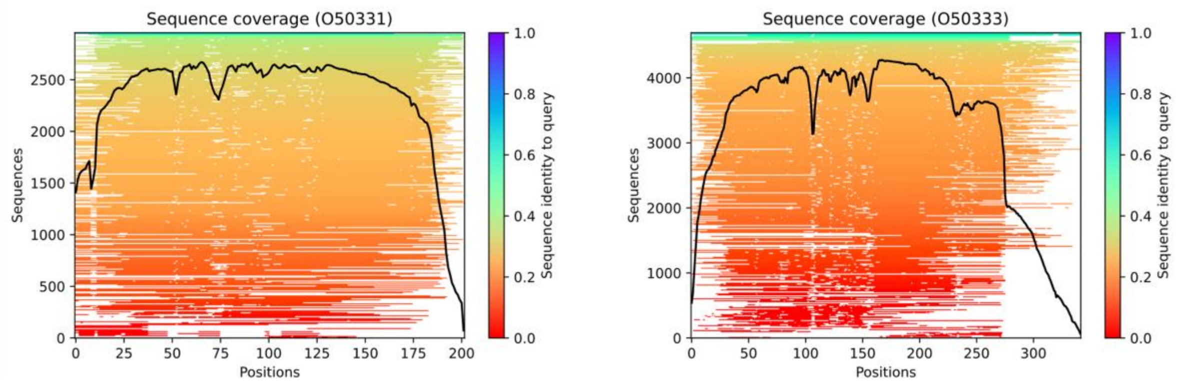


This figure illustrates the depth of the multiple sequence alignment (MSA) for O50331 (left) and O50333 (right), with sequences colour-coded by their level of identity to the reference sequence. The MSA was generated specifically for each protein analysed, providing insights into whether the alignment depth is sufficient to capture co-evolutionary signals essential for predicting both intra- and inter-protein interactions.

**SUPPLEMENTARY FIGURE 2**

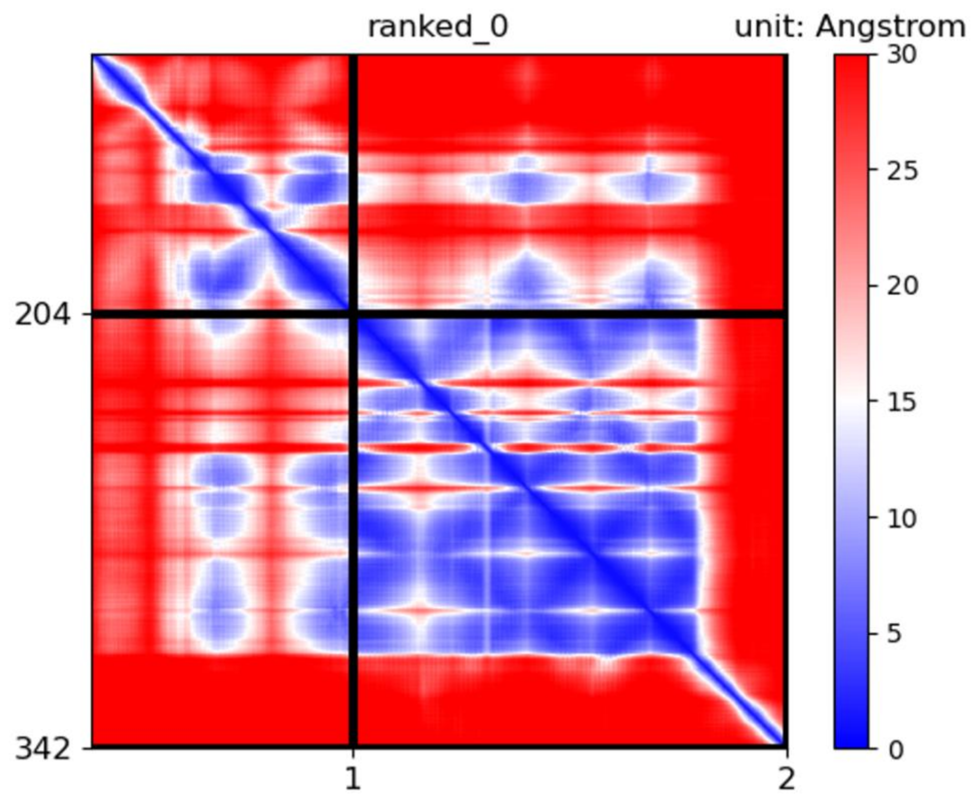


Predicted Alignment Error (PAE) plot for O50331 and O50333. This figure displays the PAE scores across the proteins, colour-coded based on the error distance in angstroms. Smaller distances indicate higher confidence in the model's predictions.

### SUPPLEMENTARY FIGURE 3

A

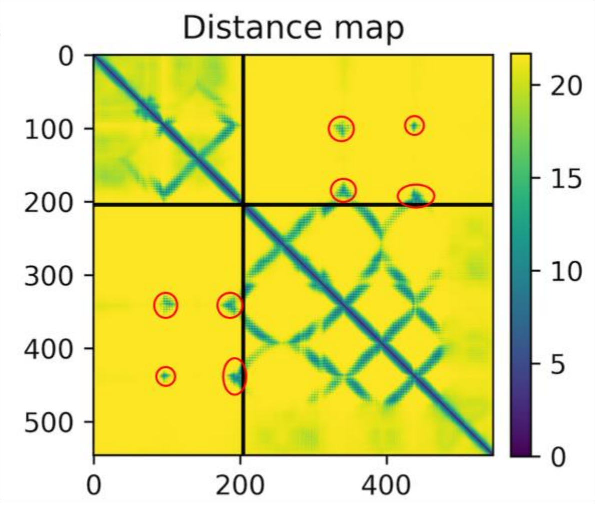


B

BC_res_int

| O50331 | O50333 | Distance Ä |
|---|---|---|
| **B:ARG 101** | C:GLU 132 | 3.4873095 |
| **B:ARG 185** | C:PHE 136 | 3.0078192 |
| **B:ARG 188** | C:GLU 133 | 2.4691393 |
| **B:ASN 189** | C:TYR 144 | 3.5194066 |
| **B:ASN 197** | C:ASN 238 | 3.3743668 |
| **B:ASN 197** | C:GLY 235 | 2.5773149 |
| **B:ASN 197** | C:GLN 237 | 3.524287 |
| **B:GLY 199** | C:ALA 236 | 3.2223024 |
| **B:GLY 199** | C:ASN 238 | 3.0896506 |
| **B:THR 202** | C:ASP 230 | 3.0321631 |
| **B:GLN 204** | C:SER 226 | 3.1155646 |

**(A)** The figure presents the distogram for O50331 and O50333, where the values indicate the protein sequence lengths in amino acids. Points near the diagonal symmetry line and within the black square represent residues in contact within the same protein, while points outside this region, highlighted by red circles, indicate contacts between the two proteins. Colours correspond to the distance between residue pairs in angstroms, with blue points indicating shorter distances. **(B)** The table provides a detailed view of the distogram, focusing on residues in direct contact between O50331 and O50333 (highlighted by red circles in the distogram). This detailed analysis offers insights into specific residue interactions within the model. The table is generated exclusively for the top-ranked protein-protein interactions that satisfy the predefined cutoff criteria.

**SUPPLEMENTARY FIGURE 4**

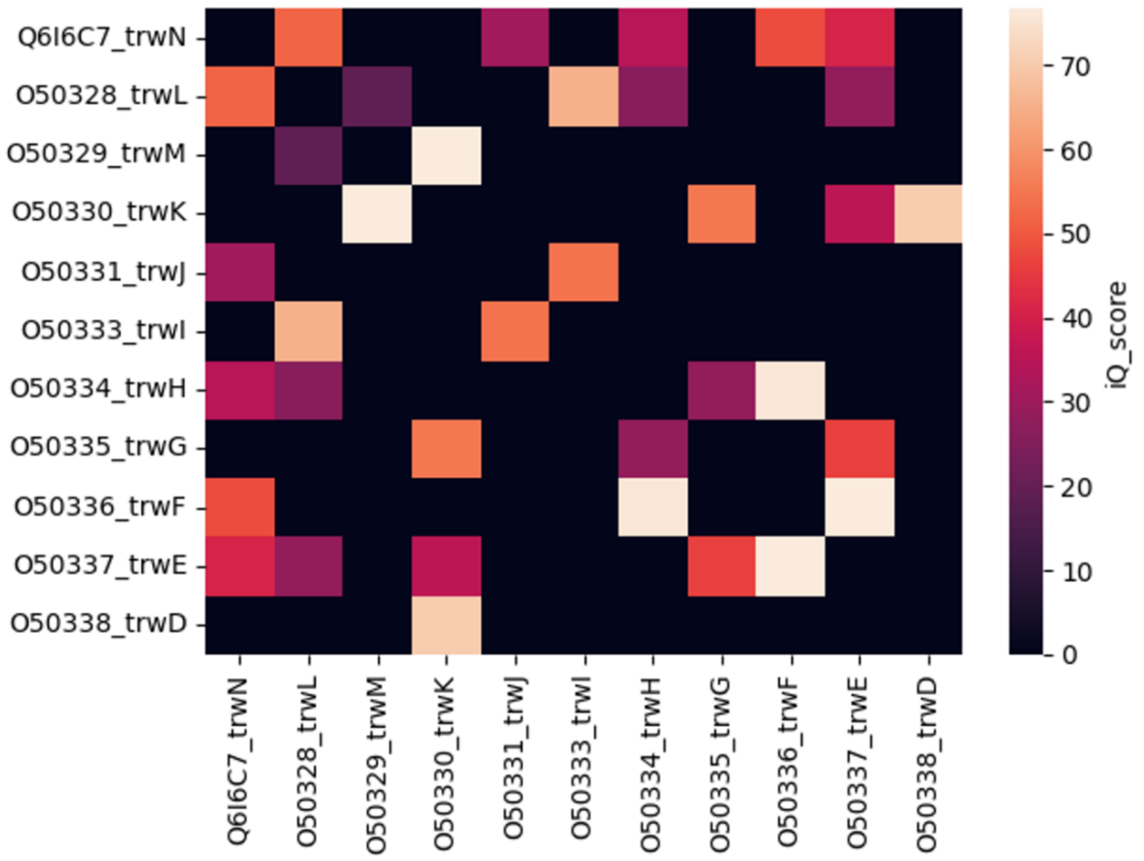


The figure displays a heatmap of interaction scores, where colours represent the iQ-scores. Higher iQ-scores are indicated by lighter colours. Black boxes denote cases with poor PAE scores, homo-oligomerisation, or excessively large total protein lengths. This matrix was constructed using data from Supplementary Data 1.

**SUPPLEMENTARY FIGURE 5**

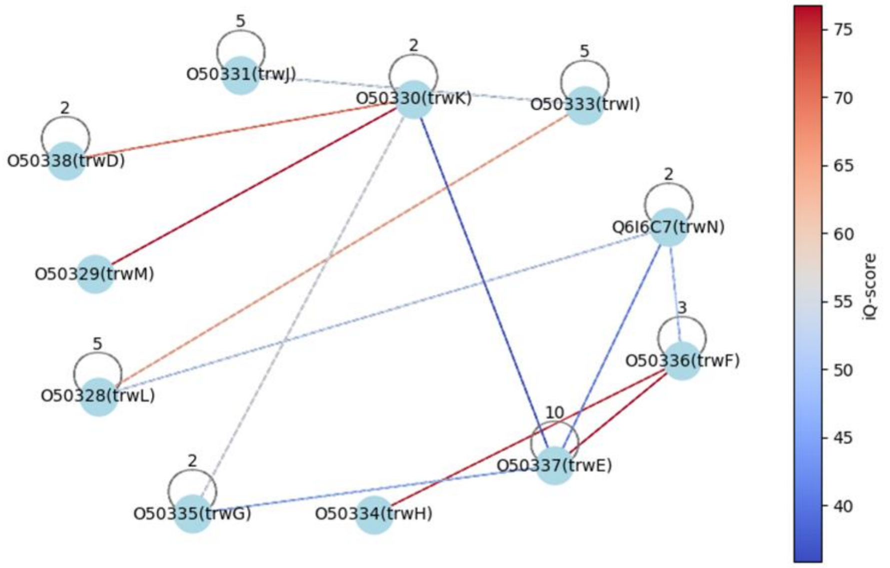


This network represents interactions within R388 proteins and is generated from Supplementary Data 1 and 2. The visualisation includes all interactions that passed the cutoff criteria, as well as homo-oligomer predictions. Each interaction is depicted as a line connecting two proteins, colour-coded according to the corresponding iQ-score. A loop on a protein indicates the most prominent homo-oligomer with the highest hiQ-score.
